# CLUE: a bioinformatic and wet-lab pipeline for multiplexed cloning of custom sgRNA libraries

**DOI:** 10.1101/2020.04.02.021279

**Authors:** Martin Becker, Heidi Noll-Puchta, Diana Amend, Florian Nolte, Christiane Fuchs, Irmela Jeremias, Christian J Braun

**Author notes:** to whom correspondence should be sent Christian J. Braun, Department of Pediatrics, Dr. von Hauner Children’s Hospital, Ludwig Maximilians University of Munich (LMU), 81377 Munich, Germany. Joint lead contact (Irmela Jeremias, Christian J Braun).

## Abstract

The systematic perturbation of genomes using CRISPR/Cas9 deciphers gene function at an unprecedented rate, depth and ease. Commercially available sgRNA libraries typically contain tens of thousands of pre-defined constructs, resulting in a complexity challenging to handle. In contrast, custom sgRNA libraries comprise gene sets of self-defined content and size, facilitating experiments under complex conditions such as *in vivo* systems. To streamline and upscale cloning of custom libraries, we present CLUE, a bioinformatic and wet-lab pipeline for the multiplexed generation of pooled sgRNA libraries. CLUE starts from lists of genes or pasted sequences provided by the user and designs a single synthetic oligonucleotide pool containing various libraries. At the core of the approach, a barcoding strategy for unique primer binding sites allows amplifying different distinct libraries from one single oligonucleotide pool. We prove the approach to be straightforward, versatile and specific, yielding uniform sgRNA distributions in all resulting libraries, virtually devoid of cross-contaminations. For *in silico* library multiplexing and design, we established an easy-to-use online platform at www.crispr-clue.de. All in all, CLUE represents a resource-saving approach to produce numerous high quality custom sgRNA libraries in parallel, which will foster their broad use across molecular biosciences.

## Introduction

Technical advances in functional genetics contribute to a growing understanding of key processes in physiology and disease. The ability to systematically perturb genomes via CRISPR/Cas9 allows assigning quantifiable phenotypes to genomic manipulations in a high-throughput format, complementing descriptive sequencing data by a functional dimension (1,2). Pooled CRISPR screens utilize the massive parallel perturbation of genes across a cell population in order to identify functional regulators of biological processes (3). Such CRISPR screens have mostly been conducted in tissue culture, but also *in vivo* (4,5).

A major limitation for performing pooled CRISPR screens is the availability of suitable sgRNA libraries (6). For example, most *in vivo* model systems do not allow studying large genome-wide libraries, as inappropriate coverage induces severe library bottlenecking, requiring small libraries, tailored to the interrogated biology of interest (7). Even for cell-line based *in vitro* screens, custom sgRNA libraries are on high demand, since they ease technical challenges, reduce costs and zoom in on a particular biological aspect of interest (8). The production of such custom sgRNA libraries, however, is technically demanding, resource-intensive and depending on certain bioinformatical expertise, which can act as an additional obstacle for many molecular biology laboratories (6).

Here, we present CLUE (<u>c</u>ustom library m<u>u</u>ltipl<u>e</u>xed cloning) – a versatile bioinformatics and wet-lab pipeline for the streamlined and multiplexed production of custom sgRNA libraries. CLUE is amenable to all major CRISPR variants including CRISPR interference (CRISPRi), CRISPR activation (CRISPRa) and CRISPR knockout (CRISPRko), covers both murine and human genomes and supports the design and production of multiplexed libraries from a single oligonucleotide pool (9). The easy-to-access web interface combines gene lists of several libraries to generate a single oligonucleotide pool. A barcoding design of unique primer binding sites enables the discrimination of individual libraries within the pool from one another and provides the foundation for the parallel generation of high quality custom sgRNA libraries. CLUE provides a straightforward, ready to use approach for cloning custom libraries, suitable for the vast majority of molecular biology laboratories.

## Results

The application of CRISPR/Cas9 for genome perturbation screens is limited by the availability of pooled sgRNA libraries. With CLUE, we set out to develop a streamlined wet-lab and easy-to-use computational pipeline for the cloning of multiple, custom sgRNA libraries from a single synthetic oligonucleotide pool.

### The concept behind the CLUE pipeline

Key to the CLUE concept is the staggered combination of three DNA adapter pairs surrounding the sgRNA sequences (Fig. 1). The outermost adapters enable PCR-based amplification of an entire synthesized oligonucleotide pool to generate double-stranded DNA, amenable for amplification and storage of the entire pool. The second adapter pair is library specific and serves as primer binding site for the specific amplification of the library of interest. Using different primer pairs allows the construction of several libraries from a single shared oligonucleotide pool. The innermost adapter pair is comprised of DNA sequences homologous to sgRNA expression vectors and amenable to Gibson cloning – therefore abolishing the need for restriction digestion of the amplified pool and enabling the inclusion of any sgRNA without any sequence restrictions.

**Figure 1:**
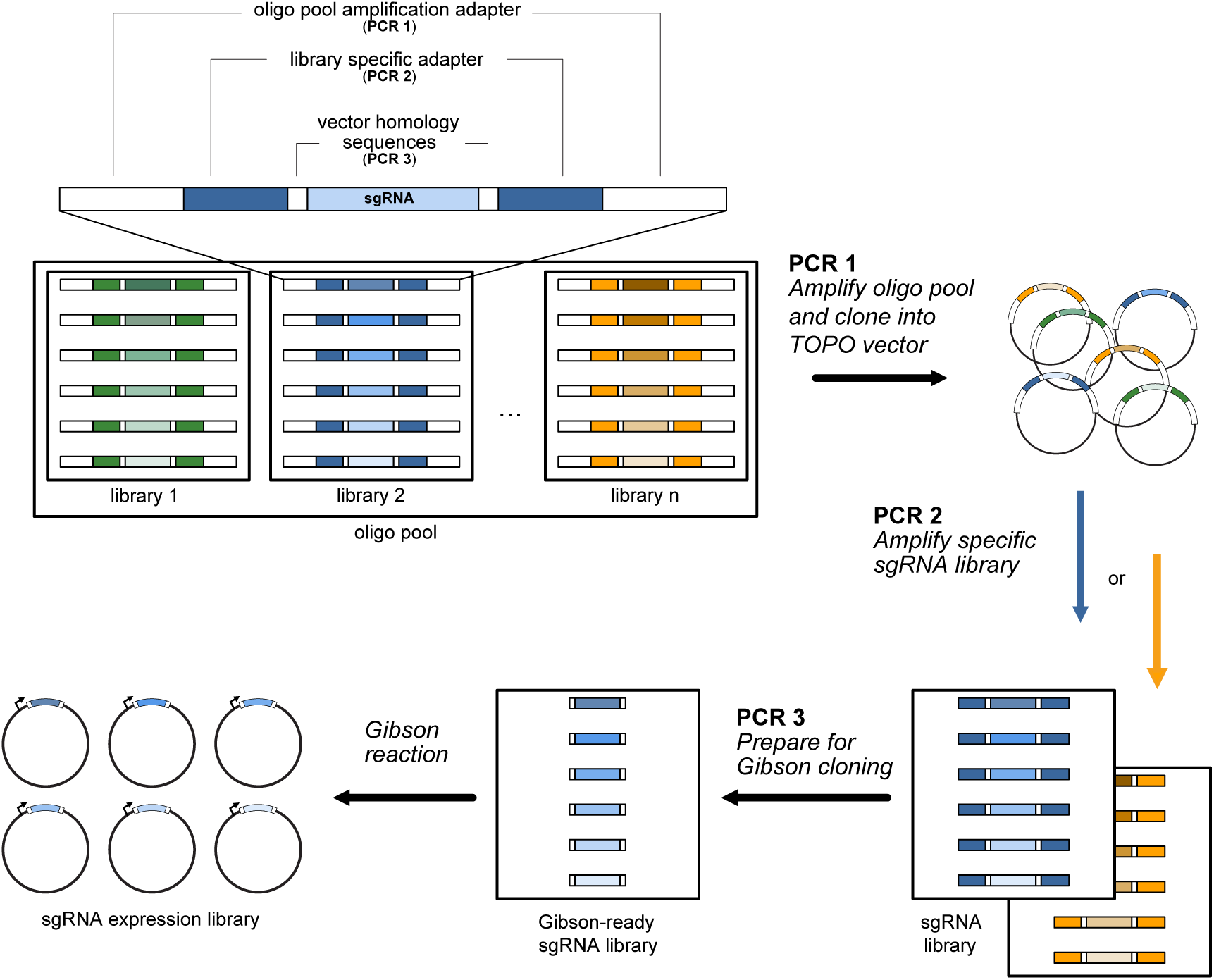
Schematic representation of the CLUE cloning pipeline. The basic structure of a CLUE oligo is the staggered combination of three DNA adapter pairs flanking a sgRNA. The outermost adapters (large white boxes, PCR 1) enable amplification of the entire oligo pool, the second adapter pair (dark colors, PCR 2) allows for specific amplification of a given sgRNA library and the third adapter pair (small white boxes, PCR 3) is comprised of sequences homologous to the sgRNA expression vector, amenable to Gibson assembly. An oligo pool comprising several sgRNA libraries is initially amplified and cloned into a TOPO vector (PCR 1). From this TOPO pool, specific libraries can be PCR amplified (PCR 2) and in a second step be prepared for Gibson cloning into a sgRNA expression vector (PCR 3) The final product is a specific sgRNA expression library.

With CLUE, library cloning requires three consecutive PCR steps (Fig. 1). After an oligonucleotide pool is obtained from a commercial vendor, the whole pool is amplified in a low-cycle-number PCR (Fig. 1, PCR1). The resulting double-stranded DNA pool is then cloned into a linearized TOPO vector, which can be amplified through transformation into competent bacteria and quality-controlled by next generation sequencing (NGS) to ensure sufficient library coverage, uniform sgRNA distribution and correct sgRNA sequences (Fig. S1A). Single distinct libraries are then amplified off the TOPO-pool through the utilization of adapter-specific PCR primers (Fig. 1, PCR2). Lastly, the library-specific PCR products are prepared for the Gibson cloning reaction into sgRNA expression vectors in a final PCR step (Fig. 1, PCR3 and Fig. S1B). The quality of the cloned sgRNA libraries can at this point be assessed by NGS.

### CLUE sgRNA libraries achieve screening-grade quality

Moving the CLUE concept to practice, we designed an oligo pool comprised of a total of 5585 sgRNAs distributed among 10 different libraries. We performed low-cycle-number amplification of the initial material (PCR1) and TOPO cloned the resulting PCR product. The re-isolated TOPO plasmid pool was subjected to NGS for quality control, showing a full sgRNA coverage of 100%. Furthermore, >90% sgRNAs were distributed within one log of the mean of the distribution (Fig. S1A). Next, we conducted library specific PCRs and the subsequent cloning steps in order to produce all 10 individual libraries (Fig. S1B), which were then subjected to NGS-based quality control. We again achieved full coverage for all libraries and never missed more than one single sgRNA per library (Table 1). Furthermore, for all libraries, sgRNAs were normally distributed with >90% of sgRNAs falling within one log of the mean of the distribution (Fig. 2 and S2, first panels). Next, we analyzed cloning and PCR accuracy and found that up to 94% of all reads matched to our initial oligonucleotide pool (Fig. 2 and S2, second panels). Lastly, we analyzed whether our cloning pipeline is indeed able to specifically amplify individual libraries without library-to-library cross contamination. For all 10 libraries cloned, we never detected >0.5% of reads mapping to other libraries than the amplified one, demonstrating high library specificity (Fig. 2 and S2, third panels). We therefore concluded that sgRNA libraries produced with the CLUE pipeline satisfy the basic quality criteria required for high-throughput pooled CRISPR screens (10).

**Table 1:** NGS-based quality assessment of 10 pooled sgRNA libraries produced with the CLUE pipeline.

| Library number | Library size (sgRNAs) | Number of sgRNAs identified by NGS | sgRNA cloning efficiency (%) | Perfectly matching NGS-reads (%) | Reads mapping to other sub-library (%) | Empty vector reads (%) | Presence of stuffer |
| --- | --- | --- | --- | --- | --- | --- | --- |
| 1 | 265 | 265 | 100.00 | 92.53 | 0.100 | 0.02 | yes |
| 2 | 705 | 705 | 100.00 | 92.64 | 0.150 | 0.01 | yes |
| 3 | 445 | 444 | 99.78 | 92.71 | 0.020 | 0.01 | yes |
| 4 | 170 | 170 | 100.00 | 93.19 | 0.010 | 0.00 | yes |
| 5 | 520 | 520 | 100.00 | 94.40 | 0.005 | 0.00 | yes |
| 6 | 530 | 530 | 100.00 | 89.68 | 0.010 | 0.00 | yes |
| 7 | 505 | 504 | 99.80 | 78.90 | 0.040 | 8.64 | no |
| 8 | 450 | 449 | 99.78 | 77.32 | 0.370 | 10.39 | no |
| 9 | 1025 | 1024 | 99.90 | 63.93 | 0.002 | 14.95 | no |
| 10 | 970 | 970 | 100.00 | 62.89 | 0.030 | 15.95 | no |

**Figure 2:**
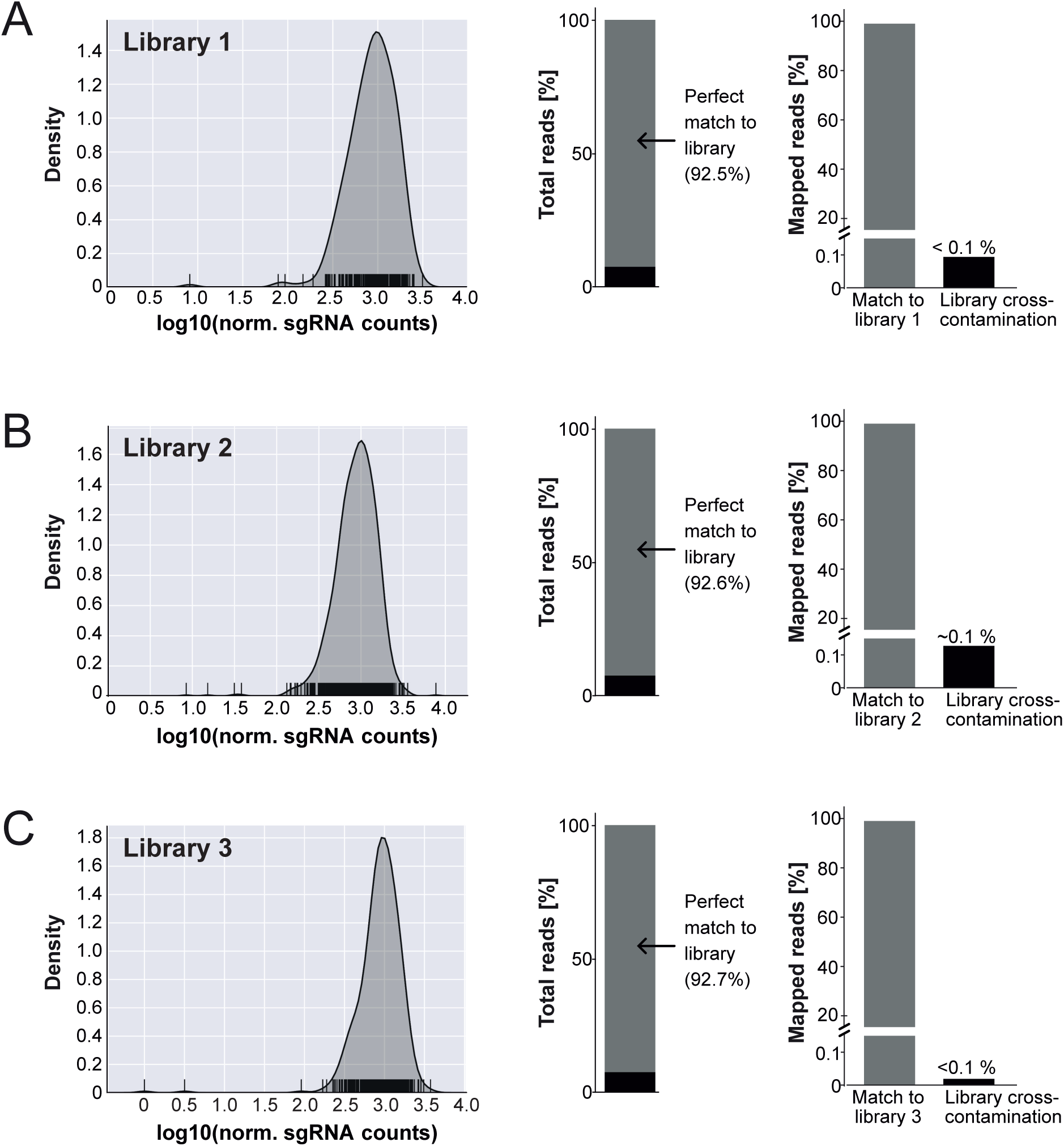
Distribution and quality assessment of three sgRNA libraries cloned with CLUE. **(A-C)** *First panel:* Density-rug plots showing the distribution of all sgRNAs of the respective library. Each rug represents one sgRNA. *Second panel:* Percentage of reads from a NGS run which could be (grey) or could not (black) be mapped to sgRNA sequences within the oligo pool. Only perfect consensus to the expected sgRNA sequences was counted as successful mapping. *Third panel:* Distribution of mapped reads from a NGS run to the library that was amplified in the given experiment (match to library) versus reads mapped to any other sgRNA library present in the original oligo pool (cross-contamination).

### A 1.2 kb stuffer reduces constructs without sgRNAs

One drawback of Gibson-based library assembly over Golden-Gate cloning is the higher prevalence of vector-only background colonies (11). Indeed, for some of the cloned sgRNA libraries we observed up to 15% of all reads mapping to expression vectors without incorporated sgRNAs (Fig. 2, S2 and Table 1). One way to enhance Gibson cloning efficacy is to improve target vector linearization. We therefore introduced a 1.2 kb stuffer sequence between the two BbsI type IIS restriction enzyme cleavage sites used for sgRNA cloning, reasoning that this would help select for perfectly linearized vector bands on agarose gels (Fig. S2, S3A, B). Indeed, this approach reduced the prevalence of empty sgRNA backbones within libraries by several hundred-fold, as demonstrated by both colony PCR and NGS (Fig. 2, S2, S3 and Table 1).

### A pilot CLUE library screen detects transcription factors essential for glioma

We next tested the capability of CLUE sgRNA libraries to produce biologically meaningful hits when employed for pooled genomic perturbation screens. To do so, we transduced the well-established human glioma cell line U87-MG with a catalytically inactive dCas9 construct in order to elicit CRISPR interference (CRISPRi). We next infected these dCas9+ U87-MG cells with a CLUE sgRNA library of 505 individual sgRNAs targeting 88 different transcription regulators (Fig. 3A). A fraction of the infected cells was kept as the input control, while the remainder was cultured *in vitro* for an additional 16 days. We extracted genomic DNA from both cohorts, amplified the integrated sgRNA loci and identified sgRNA distributions by NGS. We then scored the total library sgRNA changes and found that the majority of sgRNAs targeting genes such as the transcription factor E2F1 had depleted during glioma cell *in vitro* culture (Table S4). Intriguingly, several E2F family members including E2F1 are known to be crucial for cell cycle progression and the target of tumor suppressors such as RB (Fig. 3C) (12). We next mathematically combined the behavior of all sgRNAs targeting a specific gene in order to generate gene level scores (Fig. 3B, Table S5) (13). Besides E2F1, we found that sgRNAs targeting the Zinc finger protein 217 (ZNF217) had strongly depleted (Fig 3B, D). ZNF217 is frequently amplified in human cancer and orchestrates a wide spectrum of pro-oncogenic cellular signaling cascades (14). In summary, our pooled CRISPRi perturbation screen underlines that sgRNA libraries produced with the CLUE pipeline can highlight biologically relevant gene sets such as oncogenes.

**Figure 3:**
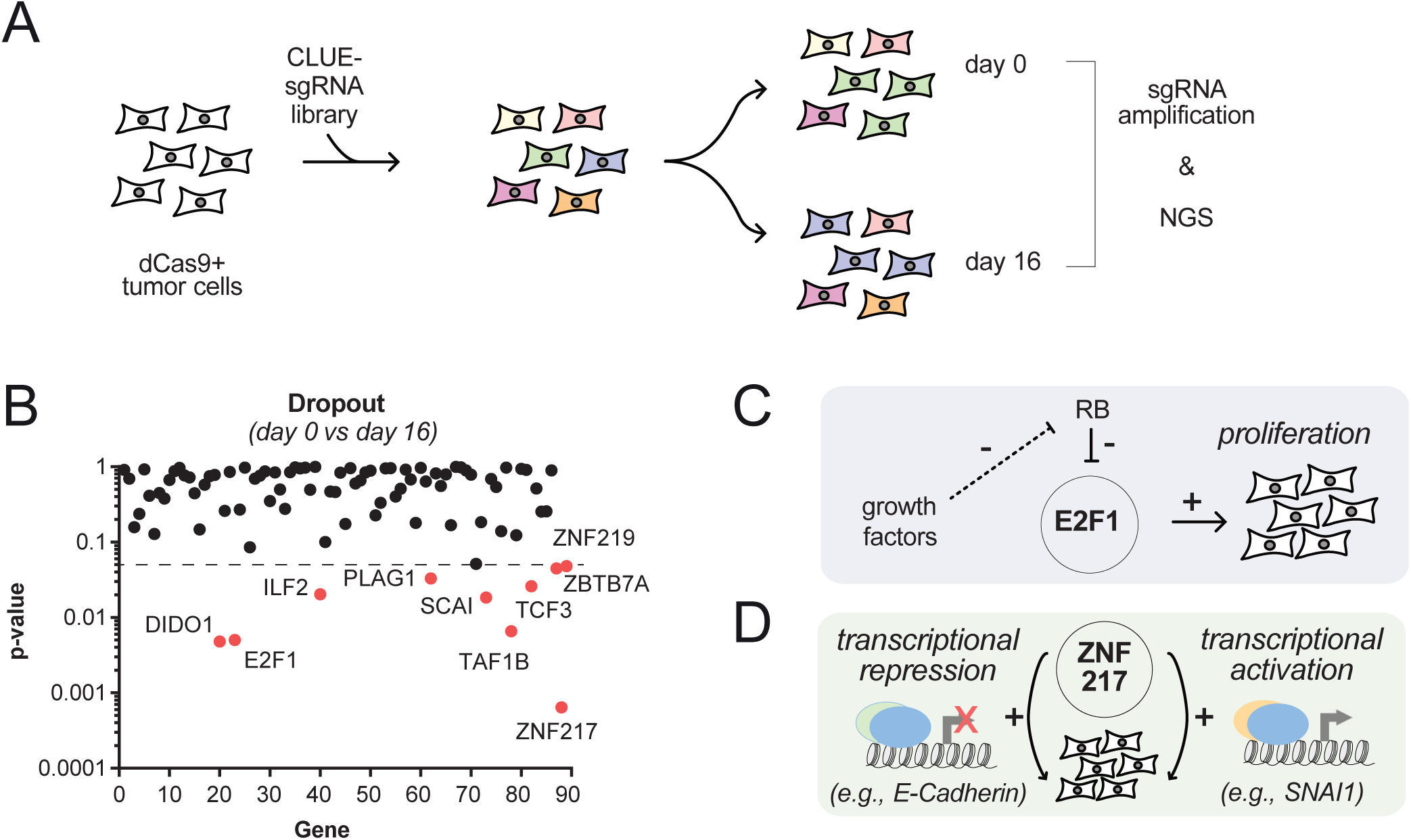
A targeted CRISPRi screen highlights transcription factors required for the proliferative fitness of malignant glioma cells. **(A)** Pooled screen schematic. U87-MG cells stably expressing a catalytically inactive dCas9 variant were lentivirally transduced with a CLUE sgRNA library of 505 individual sgRNAs targeting 88 different transcription-associated genes. **(B)** CRISPRi screen analysis. Significance of depletion after 16 days is plotted for all targeted screens. Dashed line: p=0.05. Genes depleting with p<0.05 are highlighted in red. **(C)** Schematic depicting the role of E2F1 in proliferation control. **(D)** Schematic depicting the pro-oncogenic role of ZNF217-mediated regulation of transcription.

### The CLUE web-interface streamlines multiplexed sgRNA library design

Having established that the CLUE pipeline streamlines multiplexed sgRNA library production, we next wanted to make the system broadly available to the scientific community. We felt that one major hurdle for researchers with a limited bioinformatics background could be the process of library and oligonucleotide pool design. To overcome this limitation, we generated www.crispr-clue.de, an intuitive and interactive website for fast and easy construction of sgRNA libraries and corresponding oligo pools (Fig. 4A). It allows users to upload various gene lists of interest, specify CRISPR variants (i.e. CRISPRko, CRISPRi, CRISPRa), choose between mouse and human as available model organisms, define libraries and select the number of sgRNAs per target gene. Behind the scenes, the CLUE webtool generates a multiplexed oligonucleotide pool ready to be uploaded to commercial synthesis providers, a list of adapter-binding primers for the amplification of each specific library and lists of sgRNAs distributed across the libraries. Importantly, all sgRNAs provided by CLUE reflect well-established genome-wide libraries (15-17). Alternatively, it is also possible to upload lists of sgRNA sequences directly and use the CLUE webtool for multiplexed oligonucleotide pool design. This additional layer of customization enables users to produce sgRNA libraries for any species and emerging new CRISPR applications. The upload format is a simple spreadsheet (.csv file, Fig. 4B). Additionally, we provide a detailed description and corresponding Python scripts for the quality control analysis of sgRNA libraries after NGS on the CLUE website. All in all, our easy-to-use web application aims at enabling numerous molecular biology laboratories to conduct personalized CRISPR screens, matching their unique area of interest.

**Figure 4:**
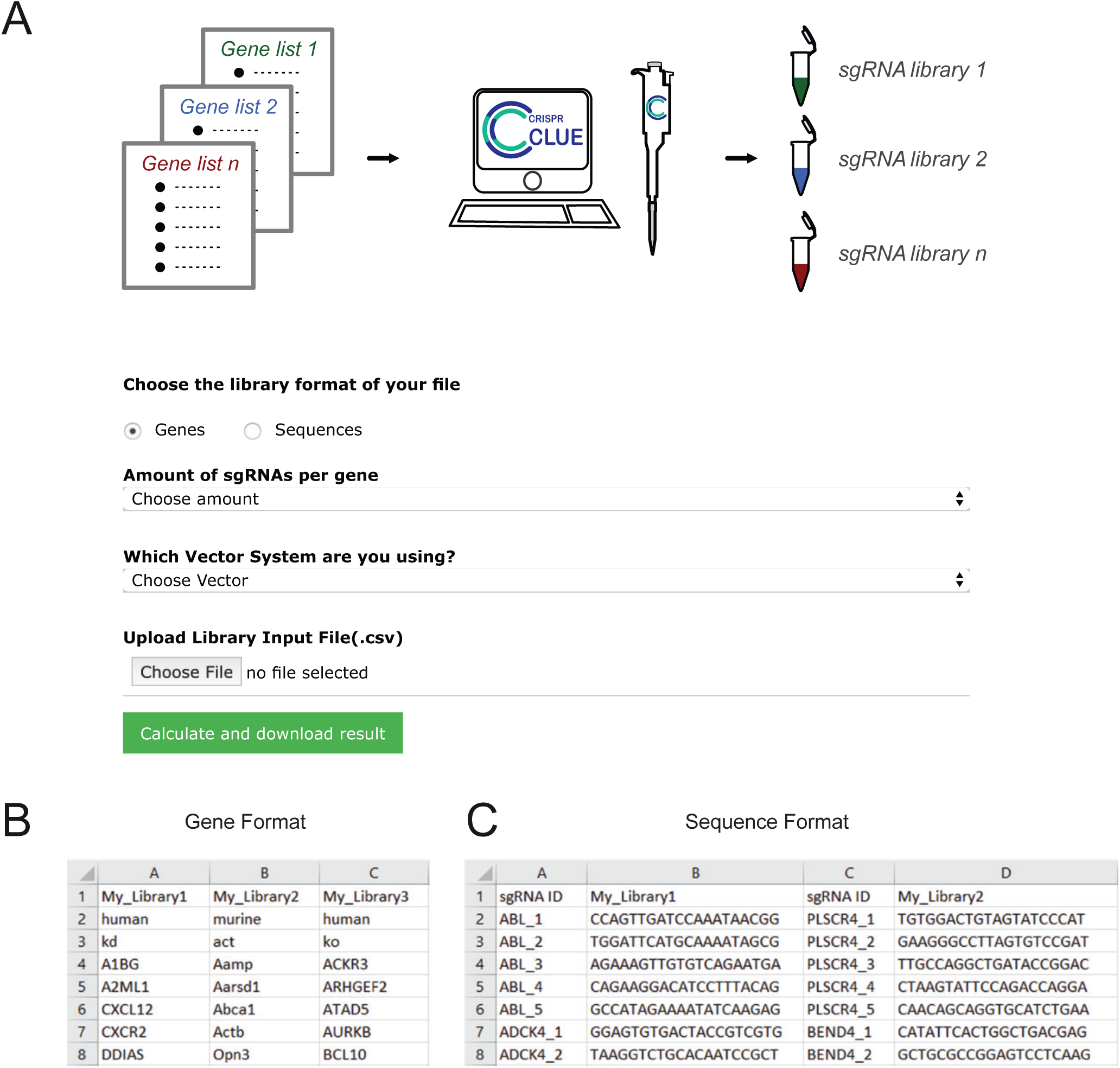
Overview of the CLUE web-interface and formats for user submission of sgRNA libraries. **(A)** Screenshot of the CLUE web-interface for the generation of oligo pools from sgRNA lists. **(B)** Example of the gene format for sgRNA libraries. Every column contains one sgRNA library, with the first cell holding the library name, the second cell holding the species (human or murine), the third cell holding the type of CRISPR application (ko, kd, act) and every following cell holding a gene name/symbol. **(C)** Example of the sequence format for sgRNA libraries. Every first column contains the string „sgRNA ID” in its first cell, followed by sgRNA IDs in subsequent cells. Every second column contains the library name in its first cell, followed by sgRNA sequences in every subsequent cell, corresponding to the given sgRNA IDs.

## Conclusion

The CLUE pipeline covers the entire workflow for multiplexed generation of custom sgRNA libraries, starting from a web-based framework for de-novo library design and multiplexing, up to a wet-lab toolset for cloning each custom sgRNA library. Libraries produced with CLUE proved to be highly accurate, with up to 94% of all NGS reads perfectly matching the intended sequences. CLUE libraries typically contain full library representation, almost no library-to-library cross-contamination and uniform sgRNA distribution within one log of the mean of the corresponding distribution – satisfying state of the art sgRNA library quality-criteria (10,18).

The CLUE pipeline is strictly optimized for efficiency. First, we incorporated an initial PCR and TOPO cloning step, which relieves constraints imposed by the limited amount of oligonucleotide pool DNA supplied by commercial vendors. Second, we utilized a barcoding protocol for library multiplexing, therefore enabling researchers to clone dozens of individual sgRNA libraries from a single oligonucleotide pool, thus reducing costs and turnaround times. Finally, the use of Gibson assembly entirely abolishes the need to avoid certain DNA sequence motifs during sgRNA library design.

We designed CLUE to be easily accessible without major skills in bioinformatics, which makes it applicable to a broad audience of biological researchers. Users can either provide lists of genes or pre-designed sgRNAs. The protocol is highly scalable, allowing individual sgRNA libraries of virtually any size.

CLUE is amenable to all major CRISPR variants including CRISPRi, CRISPRa and CRISPRko. The pipeline’s flexibility enables constructing libraries for murine and human genomes when using gene lists, but can be extended to any species of interest, when sgRNA sequences are provided. Along the same lines, this flexibility allows to extend the system to any sgRNA expression vector of interest (19). Beyond the coding genome, CLUE can be utilized for emerging new CRISPR applications such as perturbation of the non-coding genome, modification of the epigenome (20,21) as well as the Cas9-mediated editing of DNA bases (22,23). By adjusting vector-homology sequences, the pipeline holds the potential to be extended to other Cas proteins and their applications, such as Cas13-based transcriptome editing (24,25).

All in all, CLUE provides a streamlined process for generating multiple custom sgRNA libraries, for all molecular biology laboratories interested in custom genome perturbation.

## Materials and methods

### *In silico* oligo pool generation

Oligo pools are generated from input spreadsheets by text parsing methods realized in a Python script. Starting from a spreadsheet in the gene name format, each column is separated into the descriptive (first 3 cells) and the gene name part. Based on the descriptive part the reference sgRNA library is chosen based on Table S1. Next, the given number of sgRNAs/gene is selected from the reference library and a “G” is prepended if the first base is not a “G”. This step ensures efficient sgRNA transcription from the RNA Pol III promoter. For each library the sgRNA sequences are then concatenated with the adapter sequences for oligo pool amplification, specific library adapters obtained from Table S2, as well as H1 promoter and sgRNA scaffold sequences. The process is iterated over for every library, using the next library specific adapter pair from Table S2 and all oligos are finally written to the output file. Alternatively, if the sequence format is provided, sgRNA sequences are directly taken from the spreadsheet and concatenated to the adapter sequences as described above. Primer sequences are directly generated from the adapter sequences of Table S2, using the forward oligo also as primer and the reverse complement of the second oligo as reverse primer.

### Oligo Pool TOPO-Cloning

The lyophilized oligo pool was reconstituted in TE buffer at 20 ng/µl. 10 ng array synthesized oligos were used for PCR amplification with Kapa Hifi Polymerase (Roche) and primers Pool_ampl_f and Pool_ampl_r (Table S3), according to the manufacturer’s instructions. PCR setup was as follows: 98°C 3 min, 98°C 30 s, 62°C 15 s, 72°C 10 s, 72°C 2 min with a total of 15 PCR cycles (PCR1). 2 µl PCR reaction were directly taken for TOPO cloning with Zero Blunt™ TOPO™ PCR Cloning Kit (Invitrogen), according to the manufacturer’s instructions. TOPO reactions were incubated at room temperature for 30 min. Next, the reaction volume was brought up to 100 µl with water and DNA was precipitated by adding 100 µl isopropanol, 2 µl 5 M NaCl and 1 µl GlycoBlue co-precipitant (Invitrogen). Samples were vortexed and incubated at room temperature for 15 min, followed by centrifugation >16,000 x g for 15 min. Supernatants were discarded and pellets washed twice with 70% ethanol. Pellets were subsequently air-dried and reconstituted in 2 µl water. TOPO plasmids were electroporated into Endura competent cells (Lucigen) according to the manufacturer’s instructions. Bacteria were plated on LB agar containing 50 µg/ml kanamycin on 245 mm squared dishes and incubated at 37°C overnight. The next day bacteria were floated off the plates with liquid LB medium and plasmid DNA was isolated using the NucleoBond Xtra Maxi Kit (Macherey-Nagel), according to the manufacturer’s instructions. Obtained plasmid pool was quality controlled by PCR with primers M13_f and M13_r (Table S3) expecting bands of 400 bp for successfully cloned plasmids. TOPO plasmid pools were further prepared for NGS by running PCRs with P5-H1_f primers binding to the H1 portion of the cloned oligos together with P7-TOPO-5p or P7-TOPO-3p primers (Table S3), binding in the vector backbone. Two PCR reactions per sample with either of the P7-TOPO primers are required due to the random orientation of the fragments in the TOPO cloning procedure. PCRs were set up with Kapa Hifi Pol, 50 ng template and 300 nM primer each, using the following protocol: 98°C 2 min, 98°C 30 sec, 62°C 15 sec, 72°C 20 sec, 72°C 2 min with a total of 25 PCR cycles. PCR fragments were purified and submitted for NGS on an Illumina HiSeq 2000, 50 bp single-end reads aiming for >500 reads per individual oligo.

### Cloning of sgRNA libraries

To clone a specific sgRNA library included in the initial oligo pool, TOPO oligo pool was used as template for PCRs using specific primers, binding the adapters of the library of interest (PCR2). In brief, 50 pg TOPO pool were used as template together with 300 nM primer and Kapa Hifi Pol. PCR reactions were performed as follows: 98°C 2 min, 98°C 20 sec, 57 – 62°C 15 sec, 72°C 1 sec, 72°C 1 min with a total of 30 PCR cycles. Annealing temperatures were chosen depending on the adapter pair used. PCR products were purified using the NucleoSpin Gel and PCR Clean-up Kit (Macherey-Nagel) and eluted DNA was quality controlled on a 2% agarose gel (expected fragment size 105 bp). 50 pg of library specific fragment were used for PCR with primers H1_f and scaff_r to generate DNA fragments ready for Gibson cloning (PCR3). Conditions for PCR3 are identical to PCR2 and use an annealing temperature of 62°C. DNA fragments from PCR3 are purified by isopropanol precipitation as described in oligo pool cloning. Purified fragments from PCR3 are quality controlled on a 2% agarose gel (expected fragment size 65 bp, Fig. S1B). 100 ng of library fragment from PCR3 were used together with 100 ng linearized sgRNA expression vector for Gibson assembly using NEBuilder HiFi DNA Assembly Master Mix (New England Biolabs), according to the manufacturer’s instructions. After completion of the reaction, volumes were brought up to 100 µl with water and DNA was precipitated with isopropanol as described above. Pellets were resuspended in 2 µl water and subjected to electroporation into Endura competent cells (Lucigen) according to the manufacturer’s instructions. Bacteria were plated on LB agar containing 100 µg/ml ampicillin on 245 mm squared dishes and incubated at 37°C overnight. The next day bacteria were floated off the plates with liquid LB medium and plasmid DNA was isolated using the NucleoBond Xtra Maxi Kit (Macherey-Nagel), according to the manufacturer’s instructions. sgRNA libraries were further prepared for NGS by running PCRs with P5-H1_f and P7-EF1a primers. PCRs for NGS preparation were performed as described above. PCR fragments were purified and submitted for NGS on an Illumina HiSeq 2000, 50 bp single-end reads aiming for >500 reads per individual sgRNA.

### NGS data analysis for sgRNA distributions

NGS data of cloned oligo pools or sgRNA libraries were analyzed with custom Python scripts to map reads either to the entire oligo pool or a library of choice. Scripts are available in the Supplementary Material or at www.crispr-clue.de. In brief, reads from fastq files were extracted, constant parts of the sequence identified and sgRNA sequences derived thereof. Mapping required perfect matching, not allowing any mismatches. In case of mismatches, sequences were scored as unmapped.

### Pooled CRISPRi screen

U-87 MG cells (ATCC HTB-14) were a kind gift of Michael Hemann (Massachusetts Institute of Technology) and were cultured in DMEM complete medium (90% DMEM / 10% FBS). They were transduced with a lentiviral vector encoding a catalytically inactive dCas9 variant tagged with EGFP and enriched for dCas9-EGFP expressing cells by fluorescence-activated cell sorting (FACS). U-87 MG dCas9-EGFP cells were then transduced with a CLUE sgRNA library at low infection rates to minimize multiple infections per cell. Successfully transduced cells were again enriched by FACS. After a brief *in vitro* expansion, cells were either subjected to genomic DNA extraction for the t=0 days control or further passaged *in vitro. In vitro* culture conditions were kept so that at least a 250 x sgRNA representation was conserved at all times. *In vitro* samples were harvested after t=16 days. Genomic DNA was extracted using the Wizard Genomic DNA Purification Kit (Promega). sgRNA insertions were amplified from the genomic DNA using Kapa Hifi Polymerase (Roche), primers P5-H1_f and P7-EF1a_r (Table S3) and 400 ng genomic DNA per 20 µl reaction. PCR reactions were performed as follows: 98°C 2 min, 98°C 20 sec, 62°C 15 sec, 72°C 1 sec, 72°C 1 min with a total of 30 PCR cycles. PCR products were purified and sequenced on an Illumina HiSeq 2000 with 50 bp single-end reads. The NGS data was analyzed with custom Python scripts to map and count reads. Read tables were then subjected to MAGeCK analysis (13).

## Acknowledgements

CJB is supported by the Max-Eder Program of Deutsche Krebshilfe (#70113377), the Care for Rare Foundation and the Society for the Advancement of Science and Research of the LMU Medical Faculty (WiFoMed).

IJ is supported by the European Research Council Consolidator Grant 681524, a Mildred Scheel Professorship by German Cancer Aid, German Research Foundation (DFG) Collaborative Research Center 1243 “Genetic and Epigenetic Evolution of Hematopoietic Neoplasms”, DFG proposal MA 1876/13-1, Bettina Bräu Stiftung and Dr. Helmut Legerlotz Stiftung.

