## Supplement for "CLUE: a bioinformatic and wet-lab pipeline for multiplexed cloning of custom sgRNA libraries"

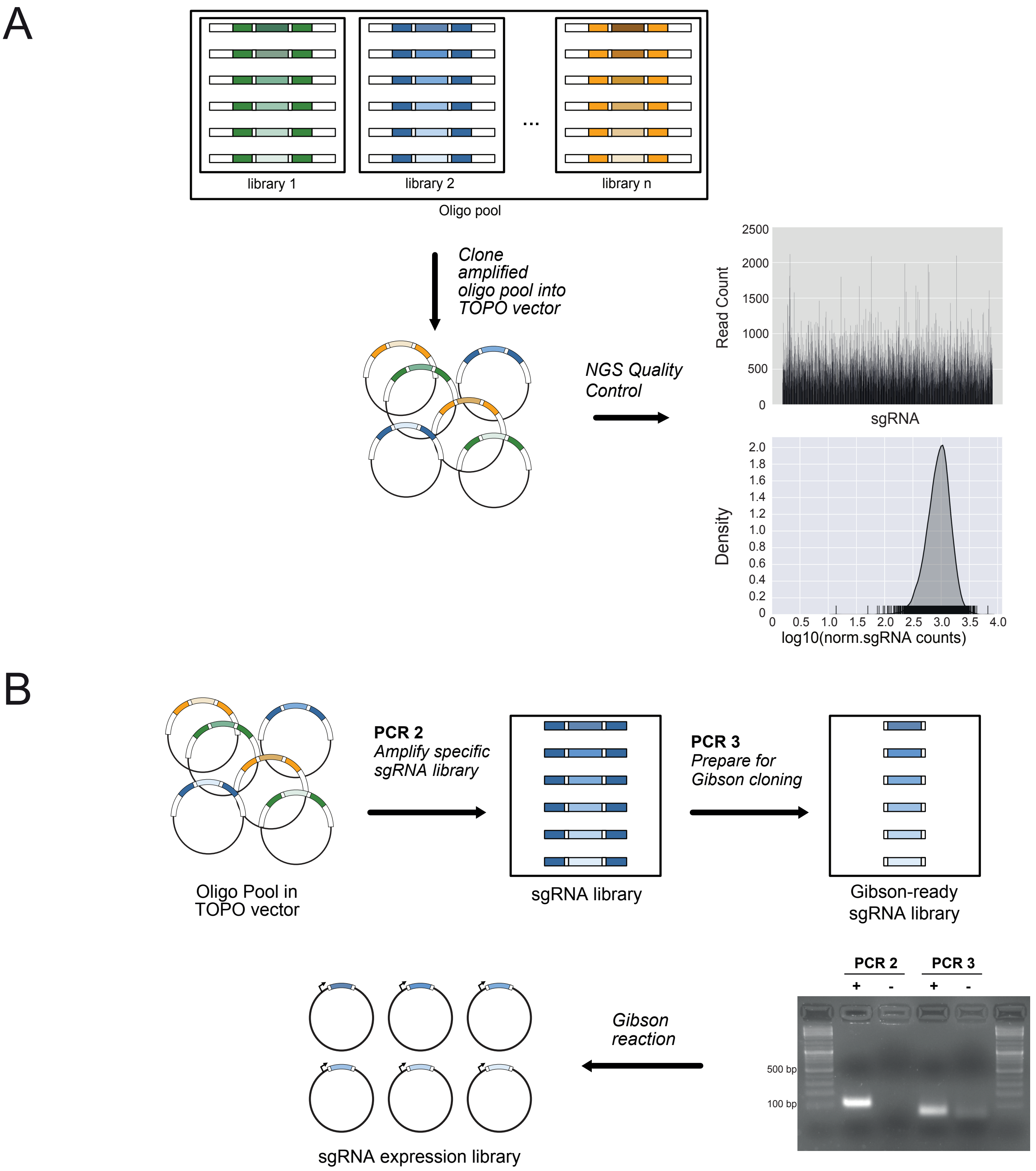


**Figure S1:** *Overview of the CLUE cloning process with quality controls.* **(A)** PCR amplification and cloning of the oligo pool into the TOPO vector is controlled by NGS. The bar graph represents raw read counts for each sgRNA of the entire pool. Normalized read counts and distribution of the sgRNAs are visualized in a density-rug plot, with each rug representing a single sgRNA. The narrowness of the distribution is a quality control measure. **(B)** PCR amplification of a given sgRNA library (PCR 2) and subsequent amplification with primers binding the vector homologous sequences to obtain smaller, Gibson-ready fragments (PCR 3), is controlled on an agarose gel, showing efficient PCR amplification and decreasing fragment size due to the nested PCR design (+ with PCR template, - water control). The Gibson-ready sgRNA library fragments are subsequently cloned into a sgRNA expression vector.

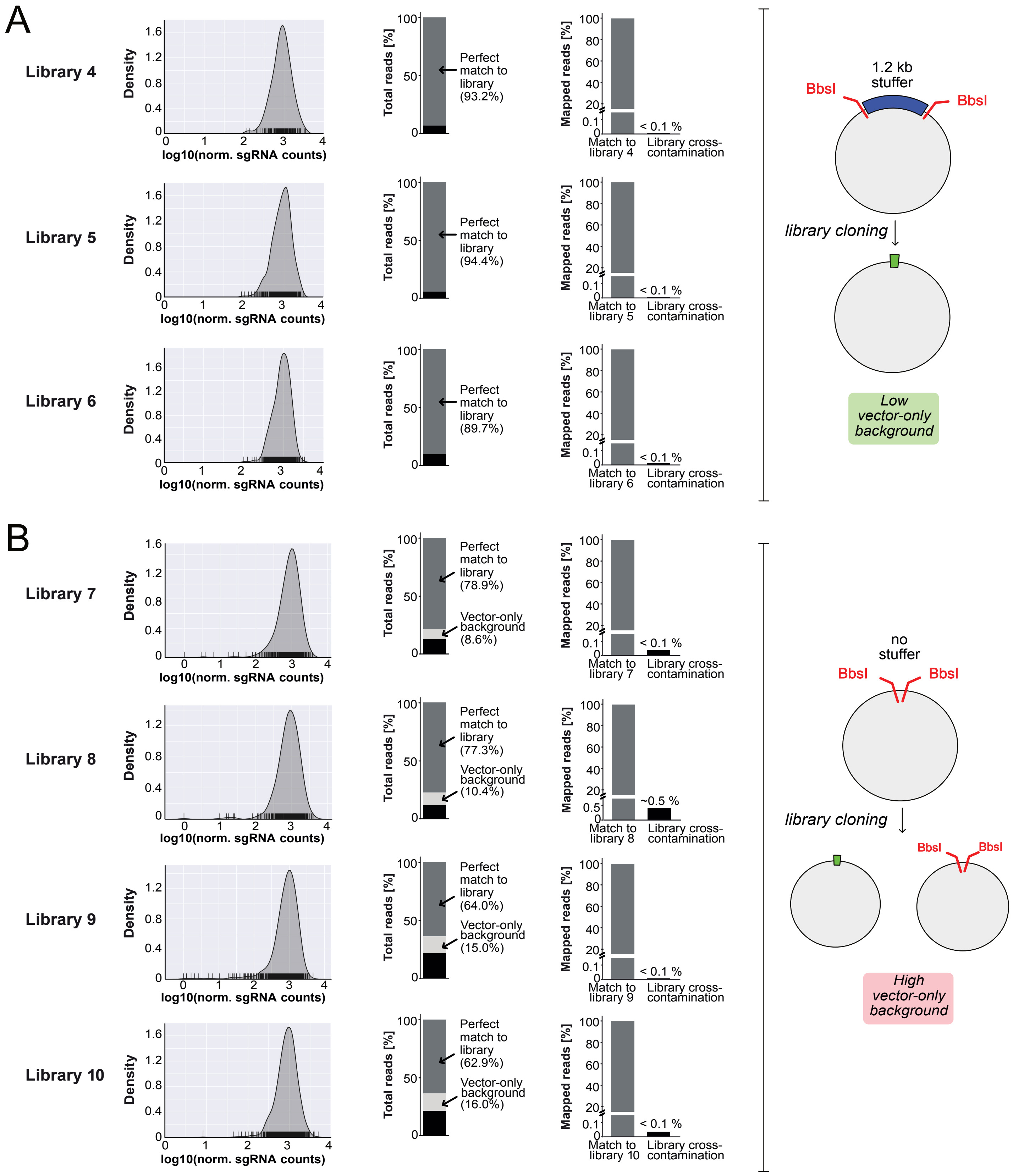


**Figure S2:** *Distribution and quality assessment of seven sgRNA libraries cloned with CLUE.* **(A)** sgRNA libraries cloned into a sgRNA expression vector containing a stuffer sequence for improved vector linearization. *First panels:* Density-rug plots revealing the distribution of all sgRNAs of the respective library. Each rug represents one sgRNA. *Second panels:* Percentage of reads from a NGS run, which could be (dark grey bar) or could not be mapped (black bar) to sgRNA sequences within the oligo pool or to the empty vector (light grey bar), respectively. Only perfect consensus to the sgRNA sequences was counted as successful mapping. *Third panels:* Distribution of mapped reads from a NGS run to the sgRNA library that was amplified in the given experiment (match to library) versus reads mapped to any other sgRNA library present in the original oligo pool (cross-contamination). **(B)** sgRNA libraries cloned into a sgRNA expression vector not containing a stuffer sequence. Panels show data as described in **(A)**.

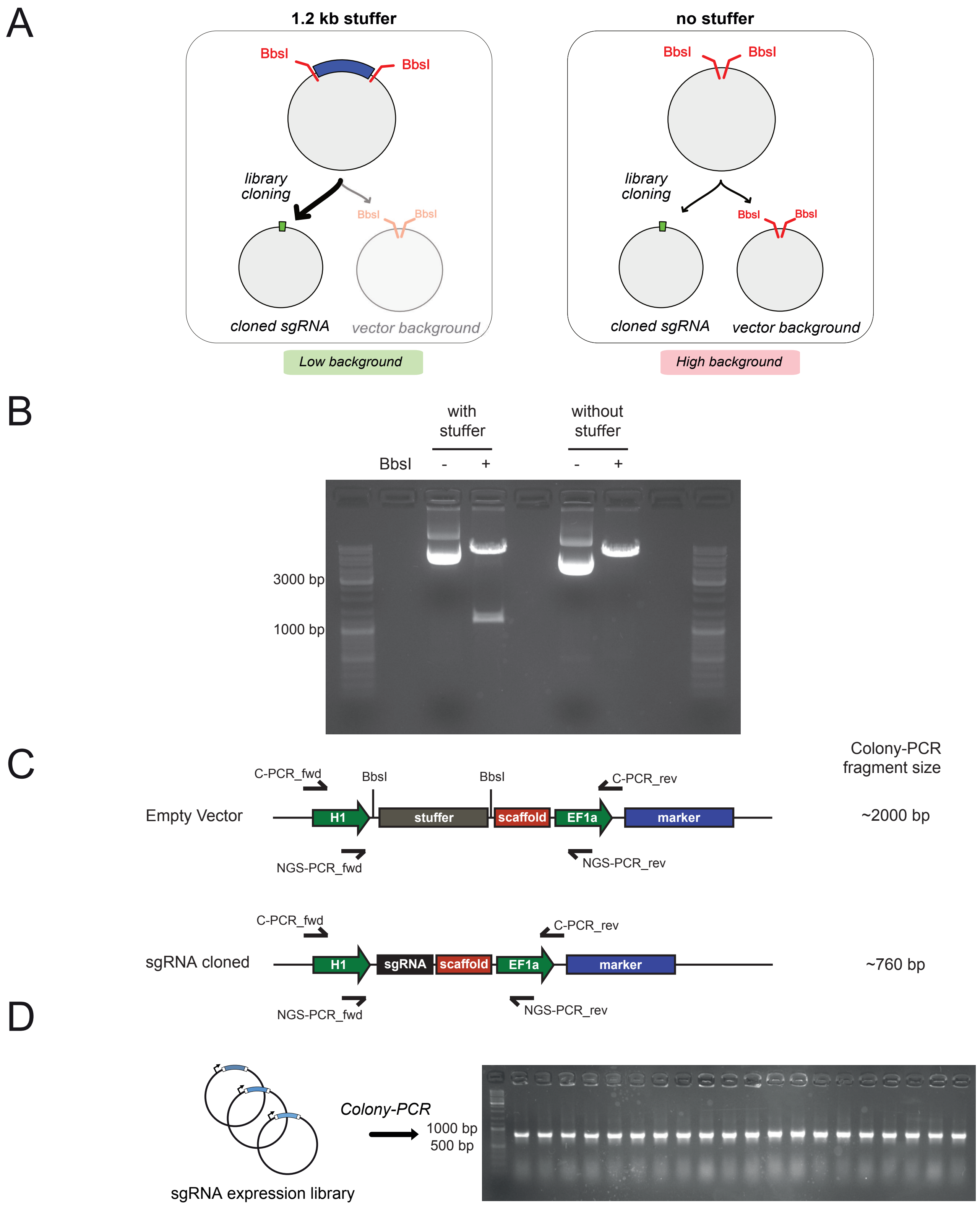


**Figure S3:** *sgRNA library cloning quality assessments.* **(A)** Schematic representation of cloning procedures using sgRNA expression vectors with or without stuffer sequences for vector linearization, respectively. Presence of a stuffer improves vector linearization and subsequent sgRNA library quality by limiting empty vector background clones. **(B)** Example agarose gel from vector linearization with BbsI, comparing the presence and absence of a stuffer sequence. **(C)** Schematic representation of sgRNA library cloning into an expression vector with stuffer sequence. Successful cloning leads to a smaller vector due to replacement of the large stuffer with a sgRNA, which can subsequently be analyzed by Colony-PCR. **(D)** Example Colony-PCR of a sgRNA library cloning experiment with an expression vector containing a stuffer sequence. All clones analyzed show a band at 760 bp, confirming successful sgRNA cloning and low empty vector background.

### Supplementary Tables

**Table S1:** Reference sgRNA libraries used by Clue

| Type | Species | Reference | Addgene ID |
|---|---|---|---|
| CRISPRko | Human | Wang et al., 2015 | #1000000067 |
| CRISPRko | Murine | Doench et al., 2016 | #73632 |
| CRISPRi | Human | Horlbeck et al., 2016 | #1000000090 |
| CRISPRi | Murine | Horlbeck et al., 2016 | #1000000092 |
| CRISPRa | Human | Horlbeck et al., 2016 | #1000000091 |
| CRISPRa | Murine | Horlbeck et al., 2016 | #1000000093 |

**Table S2:** Sequences of DNA adapters for the specific amplification of sgRNA libraries

| **Forward Adapter** | **Reverse Adapter** |
|---|---|
| TCACAACTACACCAGAAG | GCAACACTTTGACGAAGA |
| CTGTGTAATCTCCGACAC | GCCTTTGCATGTTGTGGA |
| GCGTGTTTGAATTCCACT | AAATTTCCTCGTCGGCTC |
| AGGATCTCTAGCCTCAAA | GATGAAGCATCGTAACTG |
| AACTGCGATCGCTAATGT | GTTCTCCAGTGCCTTATT |
| CTTCTACCGAACATACAG | TCGCGTTATGCTGTATGT |
| CGTAGCGTTTTGTACACG | CACGCTCAAAAAGCGTAC |
| GGCATGCTGCAATAACCT | AAACCGGTGAGCTGGAAT |
| CCGGTAACTATTCTAGCC | GCAGCCCGAATACTTTCA |
| ATACGTTAACCCGTATGG | TCTAGCCTACAATTCACG |
| CCCTAAGCATTCGCGAAA | ATTGTTAGCGCCACAATC |
| CATTTCTAGCCCTTCGAG | AACTCAACTTGGCAGGAA |
| GGATTGGACAAGCTAGTT | TAATACAAGGCCGCGTTG |
| GGGGGAAAAGGATCGATT | TCACGTGTAAACGATGCG |
| ACCCCAGTTGTGAATATC | TCGGATGACCCTAGAAAG |
| TATGAACCACTAAGGCGT | CGTAAAGTCTGCTGGTGA |
| GCTGGTGTTATGGTGGAA | AAGTCGTTTTTGGACCGC |
| GAGCACACACAAGAATGA | TAGCGACAACGTCACAAC |
| CACACTATGTCATCCGCA | CCTTTAGCAAGCAAAGCC |
| TAGTCAGAGAGTCGAGAG | GGTTTGGAGCCATTAGTT |
| GTAGACAAATGCTTGGAC | TCCCAGTCTAGTATGAGG |
| AAGGCCCAAGTCGCTTTT | CCGCCTTTTTCAAGTGAT |
| TCGATCCGGGAGTATACA | CCTACAAGAGTTCGACAC |
| ACATCCTGGTTACTTGGC | TGCTCTCGATCATAGCCT |
| GAAATATTAAGGGCGGCT | TTCGGTCAATAGAGTCGG |
| AATATGTCCCGTTCCTAC | CACTCTGTTCGGAAAACT |
| TTGGAAAGCAGTACTGCA | GATGTGTAAGTGGGCCTA |
| AGAGATCCGGTGTCAAAG | CCCAGTTCAATCGCTCAA |
| CGACTGGACGGATTTTGA | CGTTCCTCCGCCTTATTA |
| TGCGTAATGCATGTGATC | TGCTATTAAGATCCTCCC |
| ATCATCAATCACTCCCCG | TCGATCTTGAGAAGCAGT |
| GTCTATTGAAGTACCTGC | TATCTTGGAGGAGCATCG |
| TGGCATTGCTTCGTCAAG | CATGACGACTTCACCCAT |
| CCAACATGACGTTCTGTC | GAACGTCGAGTAAATGTC |
| TTTACGGTCCACCATTTG | AACGTGAGGATTAGCGCT |
| GACGTGGACTTGGACAAA | GAACTGTGCGATTTGCAG |
| TGTTTGACAATCTCGGGC | GCACATAAGTACCACTCC |
| CAAGCGCTAAGCACGAAA | AGTGATATCTACCGCGTG |
| TTATTAGTTTGTGCCCGC | AAAAGCATACAGGCACCA |
| CTCGGGATAGTTATACTC | CTTAGTGTAGATTTGGGC |

**Table S3:** List of Primers used

| Primer Name | Sequence (5' → 3') | Purpose |
|---|---|---|
| Pool_ampl_f | TGCGGATCATTCAATACGG | Initial oligo pool amplification (PCR1) |
| Pool_ampl_r | CGCCATAACGATGTTTGAG | Initial oligo pool amplification (PCR1) |
| H1_f | CTGTATGAGACCACTCTTTCCC | PCR for cloning-ready fragments (PCR3) |
| Scaff_r | TGTTTCCAGCATAGCTCTTAAAC | PCR for cloning-ready fragments (PCR3) |
| M13_f | GTAAAACGACGGCCAG | Control PCR of TOPO pool |
| M13_r | CAGGAAACAGCTATGAC | Control PCR of TOPO pool |
| P7-TOPO-5p | CAAGCAGAAGACGGCATACGAGATNNNNNNNNGTCTCGTGGGCTCGGAGATGTGTATAAGAGACAGGAGCGGATAACAATTTCACACAGG | NGS rev primer for TOPO pool |
| P7-TOPO-3p | CAAGCAGAAGACGGCATACGAGATNNNNNNNNGTCTCGTGGGCTCGGAGATGTGTATAAGAGACAGGTTTTCCCAGTCACGACGTTG | NGS rev primer for TOPO pool |
| P5-H1_f | AATGATACGGCGACCACCGAGATCTACACNNNNNNNNTCGTCGGCAGCGTCAGATGTGTATAAGAGACAG[N]$_n$GTATGAGACCACTCTTTCCCG | NGS fwd primer for TOPO pool and cloned library |
| P7-EF1a_r | CAAGCAGAAGACGGCATACGAGATNNNNNNNNGTCTCGTGGGCTCGGAGATGTGTATAAGAGACAGAAAAAGGCGGAGCCAGTACA | NGS rev primer for cloned library |
| C-PCR_f | CGATCTGCAATATTTGCATGTCGC | Colony PCR |
| C-PCR_r | TCGCTAGCTCTAGAGTAGGCGC | Colony PCR |

*[N] NGS barcodes, [N] nucleotides for staggers
